# Transmission of mechanical information by purinergic signaling

**DOI:** 10.1101/463315

**Authors:** N. Mikolajewicz, S. Sehayek, P. W. Wiseman, S. V. Komarova

## Abstract

The human skeleton constantly interacts and adapts to the physical world. We have previously reported that physiologically-relevant mechanical forces lead to small, repairable membrane injuries in bone-forming osteoblasts, resulting in the release of ATP and stimulation of purinergic (P2) calcium responses in neighbouring cells. The goal of this study was to develop a theoretical model describing injury-related ATP and ADP release, extracellular diffusion and degradation, and purinergic responses in neighboring cells. The model was validated using experimental data obtained by measuring intracellular free calcium ([Ca^2+^]_i_) elevations following mechanical stimulation of a single osteoblast. The validated single-cell injury model was then scaled to a tissue-level injury to investigate how purinergic responses communicate information about injuries with varying geometries. We found that total ATP released, peak extracellular ATP concentration and the ADP-mediated signaling component contributed complementary information regarding the mechanical stimulation event. The total amount of ATP released governed the maximal distance from the injury at which purinergic responses were stimulated, as well as the overall number of responders. The peak ATP concentration reflected the severity of an individual cell injury and determined signal propagation velocity and temporal synchrony of responses. Peak ATP concentrations also discriminated between minor and severe injuries that led to the release of similar total amounts of ATP due to differences in injury repair dynamics. The third component was ADP-mediated signaling which became relevant only in larger tissue-level injuries, and it conveyed information about the distance to the injury site and its geometry. Taken together, this study identified specific features of extracellular ATP/ADP spatiotemporal signals that encode the severity of the mechanical stimulus, the distance from the stimulus, as well as the mechano-resilient status of the tissue.

## 2 Introduction

The human skeleton is constantly interacting and adapting to the physical world, as seen by the loss of bone in astronauts experiencing microgravity (1) or gain of bone in athletes engaged in intense activities (2). The magnitude, frequency and duration of mechanical loading are known determinants of the mechano-adaptive response of bone at the tissue-level (3-5) and at the cellular-level, as observed in intracellular free calcium ([Ca^2+^]_i_) elevations of osteocytes during *in vivo* mechanical loading (6).

We have recently reported that physiologically-relevant mechanical loading routinely injured bone cells *in vitro* and *in vivo*, resulting in release of ATP through plasma membrane disruptions, and stimulation of calcium responses in the neighbouring cells (7). These membrane disruptions in bone cells are counteracted by rapid vesicle-mediated membrane repair (7, 8), which limits ATP spillage. Thus, contrary to previous generalizations that ATP is released as a bolus proportional to mechanical stimulus (9), our data suggest that mechanically-stimulated ATP release contains dynamic information about both the extent of the injury and the rate of repair.

ATP stimulates autocrine and paracrine [Ca^2+^]_i_ signaling through purinergic (P2) receptor network, which consists of seven ionotropic P2X receptors and eight G-protein coupled P2Y receptors (10-12). Several P2 receptors have been implicated in the mechano-adaptive response in bone (13-15), however it is uncertain why so many receptors are required and how they integrate mechanical information. While individual P2 receptors are sensitive to ATP concentrations over 2-3 orders of magnitude, the entire P2 receptor network covers over a million-fold range of ATP concentrations (16, 17). In addition, many P2Y receptors are differentially sensitive to ADP and ATP. Thus, it is possible that unique subsets of P2 receptors are activated in neighboring bone cells depending on the position and severity of the mechanical stimulus.

We hypothesized that mechanically-stimulated ATP release dynamics reflect the balance between cell membrane injury and repair; and that ATP and ADP released from the injury site generate unique spatiotemporal signatures that convey information to non-stimulated neighboring cells about the mechanical stimulus, such as severity and distance to stimulus, as well as the state of tissue mechano-resilience. We developed a mathematical model to account for injury-related ATP and ADP release, their extracellular diffusion and degradation, as well as paracrine purinergic responsiveness of neighboring cells. The experimentally validated model was scaled to a tissue-level mechanical stimulus to investigate how a cellular population responds to mechanical stimuli with position- and severity-appropriate responses.

## 3 Materials & Methods

### Solutions

Phosphate-buffered saline (PBS; 140 mM NaCl, 3 mM KCl, 10 mM Na_2_HPO_4_, 2 mM KH_2_PO_4_, pH 7.4), sterilized by autoclave. Physiological solution (PS; 130 mM NaCl; 5 mM KCl; 1 mM MgCl_2_; 1 mM CaCl_2_; 10 mM glucose; 20 mM HEPES, pH 7.6), sterilized by 0.22 μM filtration. Bioluminescence reaction buffer (0.1 M DTT, 25 mM tricine, 5 mM MgSO_4_, 0.1 mM EDTA, 0.1 mM NaN_3_, pH 7.8), sterilized by 0.22 μm filtration.

### Reagents

Minimum Essential Medium (MEM) α (Gibco 12,000-022); Fura2 AM (Invitrogen F1221); D-Luciferin potassium salt (Invitrogen L2916); Collagenase P from *Clostridium histolyticum* (Roche 11213857001); adenosine 5’-triphosphate magnesium salt (Sigma-Aldrich A9187), adenosine 5’-diphosphate sodium salt (Sigma-Aldrich A2754), adensoine 5’-monophosphate disodium salt (Sigma-Aldrich 01930), adenosine (Sigma-Aldrich A4036), hexokinase from *Saccharomyces cerevisiae* (Sigma-Aldrich H6380), L-ascorbic acid 2-phosphate sesquimagnesium salt hydrate (Sigma-Aldrich A8960), luciferase from *Photinus pyralis* (Sigma-Aldrich L9420), phosphoenolpyruvic acid monopotassium salt (Sigma-Aldrich 860077), pyruvate kinase from rabbit muscle (Sigma-Aldrich P9136), quinacrine dihydrochloride (Sigma-Aldrich Q3251); Venor GeM Mycoplasma PCR-based detection kit (Sigma-Aldrich MP0025); ARL 67156 trisodium salt (Tocris Bioscience 1283) and Suramin hexasodium salt (Tocris Bioscience 1472); Dulbecco’s Modified Eagle Medium (Wisent Bio Products 319-020 CL), fetal bovine Serum (Wisent Bio Products 080152); Penicillin Streptomycin (Wisent Bio Products 450-201-EL); sodium pyruvate (Wisent Bio Products 600-110-UL); collagenase Type II (Worthington Biochemical Corporation LS004176).

### Cell Culture

All procedures were approved by McGill’s University’s Animal Care Committee and complied with the ethical guidelines of the Canadian Council on Animal Care. See Supplementary Materials for detailed methodology. For compact bone-derived osteoblasts (CB-OB), 4-6 week old C57BL/6 mice (Charles River) femurs and tibia bone fragments were enzymatically digested and cultured for 3-5 days in αMEM (supplemented with 10% FBS, 1% sodium pyruvate, 1% penicillin streptomycin) as described previously (7). Cells were trypsinized, filtered, plated at 10^4^ cells/cm^2^ in osteoblast differentiation medium (+ 50 μg/mL ascorbic acid) and cultured for 2-3 days prior to experiments. The C2C12 cell line (ATCC CRL-1772), stably transfected with BMP-2 (C2-OB, courtesy of Dr. M. Murshed, McGill University) to acquire osteoblastic phenotype, was plated at 10^4^ cells/cm^2^ in DMEM (supplemented with 1% penicillin streptomycin, 1% sodium pyruvate and 10% FBS) and cultured for 2-3 days prior to experiments. Absence of mycoplasma contamination was verified in cryo-preserved stocks of C2-OB cells using PCR-based detection kit.

### Intracellular calcium recordings and analysis

Cells plated in glass-bottom 35 mm dishes or 48-well plates (MatTek Corporation) were loaded with Fura2-AM and imaged in physiological solution as previously described (18). The data were characterized using a previously developed MATLAB algorithm (19).

### ATP/ADP solutions

Purine nucleotide solutions were enzymatically treated to deplete undesirable purine nucleotide species. ADP present in ATP solutions was converted to ATP by 20 U/mL pyruvate kinase in the presence of excess phosphoenolpyruvate. ATP present in ADP solutions was converted to ADP by 20 U/mL hexokinase. All enzymatic reactions were prepared in physiological solution and incubated at 37°C for 30 min followed by 2 min heat inactivation at 95°C. Solutions were always prepared on the day of experiment.

### Single cell mechanical stimulation

Cells were acclimatized for 10 min on the stage of an inverted fluorescence microscope (Nikon T2000). A micropipette was positioned approximately 10 μm from the cell membrane at a 45° angle from the horizontal plane and mechanically stimulated at a speed of 250 μm/s with a contact duration of 60 ms using a FemtoJet microinjector NI2 (Eppendorf Inc.).

### Vesicle release kinetics

Cells were incubated with 10 μM quinacrine solution for 15 min at room temperature and washed with PBS. Time-lapse recordings of quinacrine-loaded cells were acquired with a Nikon T2000 inverted fluorescence microscope at a sampling rate of 2 Hz. Vesicular release was identified as events of sudden loss of localized fluorescence using an ImageJ plugin SparkSpotter (Babraham Bioinformatics, UK) applied to temporally reversed and contrast-enhanced image stacks.

### ATP measurement

100 μL of supernatant was collected and ATP content was measured by bioluminescence luciferin-luciferase assay using FB 12 single tube luminometer (Titertek-Berthold).

### Statistical analysis

Data are representative images and traces, or means ± S.E.M, with *N* indicating the number of independent trials. Statistical significance was assessed by ANOVA followed by a Bonferroni post-hoc test, results were accepted as significant at p<0.05. Numerical simulation of diffusion profiles was conducted in Mathematica 11.2 (Wolfram). Curve fitting and model-related analyses was conducted in MATLAB R2018a (MathWorks). Logistic regression and ROC curves were generated in SPSS 24 (IBM).

During analysis of numerical simulations, the parameter space that predicted outcomes *θ_pred_* most consistent with experimental observations *θ_obs_* was identified by computing the *Bias* between *θ_pred_* and *θ_obs_* as the root difference between mean squared error (MSE) and variance (Var),

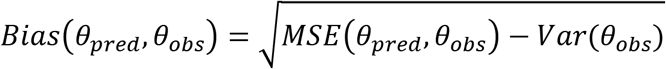
Where

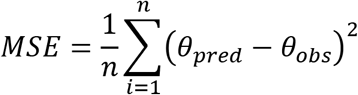

When two sets of outcome measurements were available (i.e., response times and response probabilities), two sets of bias measurements *Bias*_1_ and *Bias*_2_ were combined as,

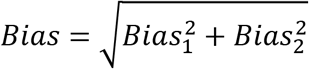
and the parameter space with minimal *Bias* was taken as the best-fit parameter space.

## 4 Mathematical Model

### ATP release model

To model ATP and ADP release from a mechanically-stimulated cell, we assumed the following:

1. cellular contents are released through a circular membrane-injury with diameter *d*(*t*) (μm), which reseals exponentially from an initial diameter *d*_0_ with a characteristic repair half-time of *τ*_1/2_ (*s*):

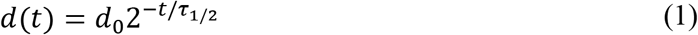
2. The rate of release of an intracellular nucleotide (ATP or ADP) can be tracked as the difference between its initial concentration *c*_0_ and current concentration *c_in_*, the rate of change which is proportional to diffusion-dependent transport coefficient *β* (*s*^−1^):

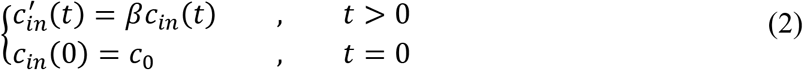
where ′ indicates a time derivative.
3. Transport coefficient *β* (*s*^−1^) depended on the injury diameter *d*(*t*), cell volume *V_cell_* (estimated to be 2400 μm^3^ for C2-OB cells), and diffusion of ATP/ADP within the cell to the site of injury at the membrane with an intracellular diffusion coefficient *D_cyto_* [*D_cyto_* ≅ 0.25*D* (20)], and then diffusion away from the injury into the extracellular space with the diffusion coefficient *D*:

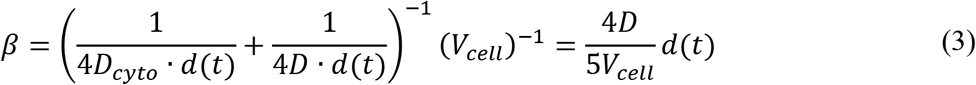

Model parameters are provided in **Table 1**.

**Table 1.** Model Parameters

| Symbol | Definition | Value(s) | Reference |
| --- | --- | --- | --- |
| **Injury, diffusion and degradation model** |  |  |  |
| $c_{0,ATP}$ | Initial ATP concentration | 5 mM | (23) |
| $c_{0,ADP}$ | Initial ADP concentration | 0.5 mM | (23) |
| $\tau_{1/2}$ | Reseal half-time constant | 0.1 to 20 s | Estimated in study |
| $d_0$ | Initial injury (diameter) | $10^{-4}$ to $10^1 \mu\text{m}$ | Estimated in study |
| $V_{\text{cell}}$ | Osteoblast cell volume | $2400 \mu\text{m}^3$ | Estimated for C2-OB cells |
| $D_{ATP}$ | ATP diffusion coefficient | $347 \mu\text{m}^2\text{s}^{-1}$ | (24-26) |
| $D_{ADP}$ | ADP diffusion coefficient | $377 \mu\text{m}^2\text{s}^{-1}$ | |
| $k_{ATP}$ | ATP degradation constant | $0.05 \text{ min}^{-1}$ | Estimated in study |
| $k_{ADP}$ | ADP degradation constant | $0.25 k_{ATP}$ | Approximated from (27) |

### ATP diffusion and degradation model

To account for geometrical constraints imposed by our *in vitro* experiments, we modelled the mechanically-stimulated cell as a finite sphere (defined by radius *R_cell_*) with uniform intracellular ATP and ADP concentrations 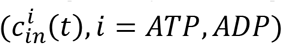 located on a flat surface. To calculate extracellular ATP and ADP concentrations (*c_i_*(*x*, *t*), *i* = *ATP*, *ADP*), we assumed the 3D radial diffusion equation with spherical boundaries and extended the problem symmetrically for negative values for ease of computation. Mathematically, this is equivalent to doubling the concentrations in a half-sphere, while setting the concentrations in the other half to 0 for a no-flux (Neumann) boundary condition, which we imposed to conserve particle number within the boundaries. To account for ATP and ADP extracellular degradation, we assumed the first-order degradation rate constant of ATP to ADP as *k_ATP_*, and the first-order degradation rate constant of ADP out of the system as *k_ADP_*. The resulting equations for ATP (Eq. 4) and ADP (Eq. 5) are given below:

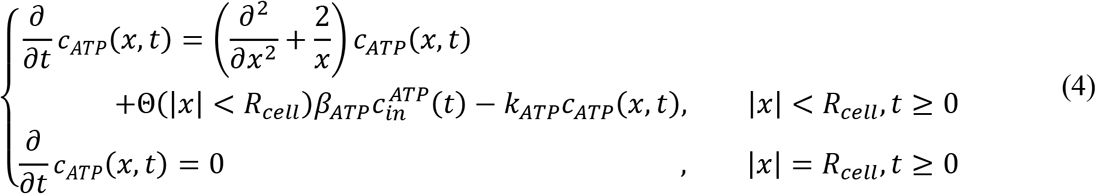

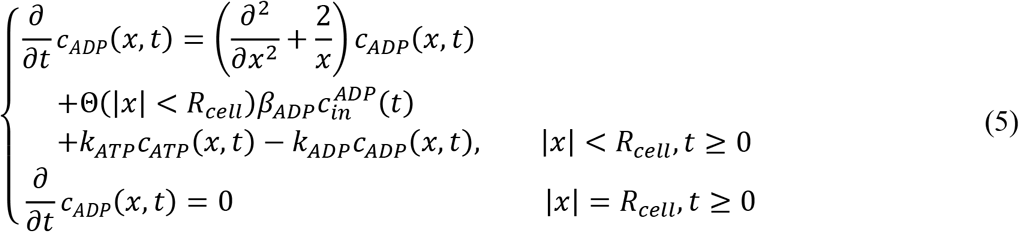
where, Θ(∙) denotes the indicator function (1 when true, otherwise 0). Eq. 4 and 5 were solved using the NDSolve routine provided by Mathematica 11.2. To avoid the singularity in the Laplacian written in spherical coordinates, the ‘MethodOfLines’ option was used with a spatial grid containing an even number of points. Model parameters are provided in **Table 1**.

### Modeling spatiotemporal response probabilities

To model the spatiotemporal paracrine responses following mechanical stimulation of an individual cell, a response time model (21) was adopted. We assumed that at location *x* from the origin and at time *t* following mechanical stimulation, all cells can be divided into three groups: (i) cells that have responded to the stimulus (*R*), (ii) cells that have not yet responded, but will respond eventually (*S*_1_) and (iii) cells that will not respond to the stimulus (*S*_0_). If *R*, *S*_1_ and *S*_0_ indicate proportions of cells, then

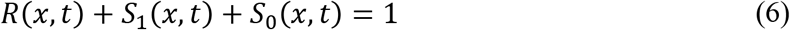

Initially, all cells are in *S*_0_ state, *S*_0_(*x*, 0) = 1. When nucleotides arrive, they stimulate a proportion of cells to respond with probability *P* (i.e., effective response probability), which is dependent on ATP and ADP concentrations. *P*(*x*, *t*) was obtained from instantaneous response probabilities *p* established from experimental dose-dependencies for ATP and ADP, spatiotemporal concentrations of which were simulated using the diffusion/degradation model (Eq. 4, 5). *P* was then calculated as the time-cumulative maxima of *p* at a given distance:

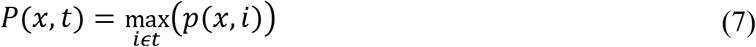

Thus, the effective response probability *P* increases over time and is related to the proportion of responding (*R*) and eventually responding (*S*_1_) cells by *P*(*x*, *t*) = *R*(*x*, *t*)+*S*_1_(*x*, *t*). Simultaneously, the proportion of cells that will never respond decreases so that*, S*_0_(*x*, *t*) = 1 − *P*(*x*, *t*).

Upon stimulation of cells with nucleotides, cells do not respond instantaneously. Instead, there is a delay time in the response which is dependent on nucleotide concentrations. We introduced *t_rxn_*(*x*, *t*) as the reaction time of purine-mediated responses obtained from experimental dose-dependencies for ATP and ADP, spatiotemporal concentrations of which were determined from the diffusion/degradation model (Eq. 4, 5). We assumed that instantaneous rate of response *λ*(*x*, *t*) is the reciprocal of the reaction time:

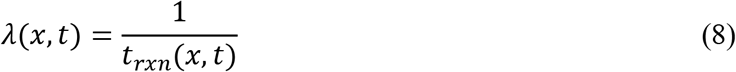
which allowed us to calculate the time-dependent proportion of cells that have not responded yet in the eventually responsive component *S*_1_:

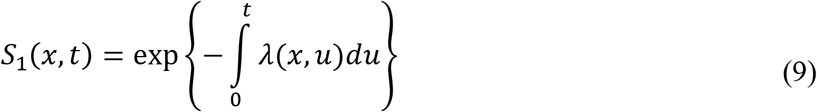

To estimate the expected distance-dependent response time *T*(*x*), we accounted for the presence of non-responding cells *S*_0_ using a conditional survival function *S*^*^:

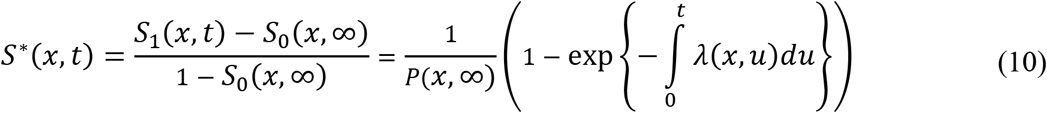
which was integrated to obtain the expected response time *T*(*x*) at distance *x*:

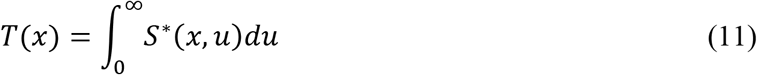

The spatiotemporal distribution of paracrine responses was then described by the probability density function *f* corresponding to *S*^*^ (22),

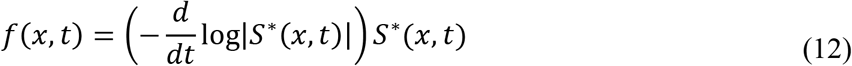

## 5 Results

### 5.1 Mechanical stimulation of a single osteoblast leads to release of purinergic signals that convey position- and magnitude-related information to neighboring osteoblasts

BMP-2-transfected C2C12 osteoblast-like cells (C2-OB) and compact bone-derived primary osteoblasts (CB-OB) were loaded with calcium-indicator Fura2 and a single osteoblast was stimulated with a glass micropipette. Shortly after, non-stimulated neighbouring cells exhibited elevations in intracellular free calcium concentrations ([Ca^2+^]_i_) which were inhibited by the purinergic antagonist suramin (**Fig 1A**). The magnitudes of these secondary responses were proportional to the magnitude of [Ca^2+^]_i_ elevation in the mechanically-stimulated primary cell (**Fig 1B**). The response probability of neighbouring cells decreased proportionally to the distance from the mechanically-stimulated cell (**Fig 1C**), while the delay to the onset of response increased (**Fig 1D**). Using a logistic regression model, we demonstrated that the characteristic parameters of the primary cell Ca^2+^ response (PR) and distance to the primary cell (dist.) predicted the onset of secondary responses with 76-79% accuracy (p<0.001, **Fig 1E**). Thus, our data suggest that purinergic signals convey information regarding the magnitude and position of the mechanical stimulus.

**Figure 1.**
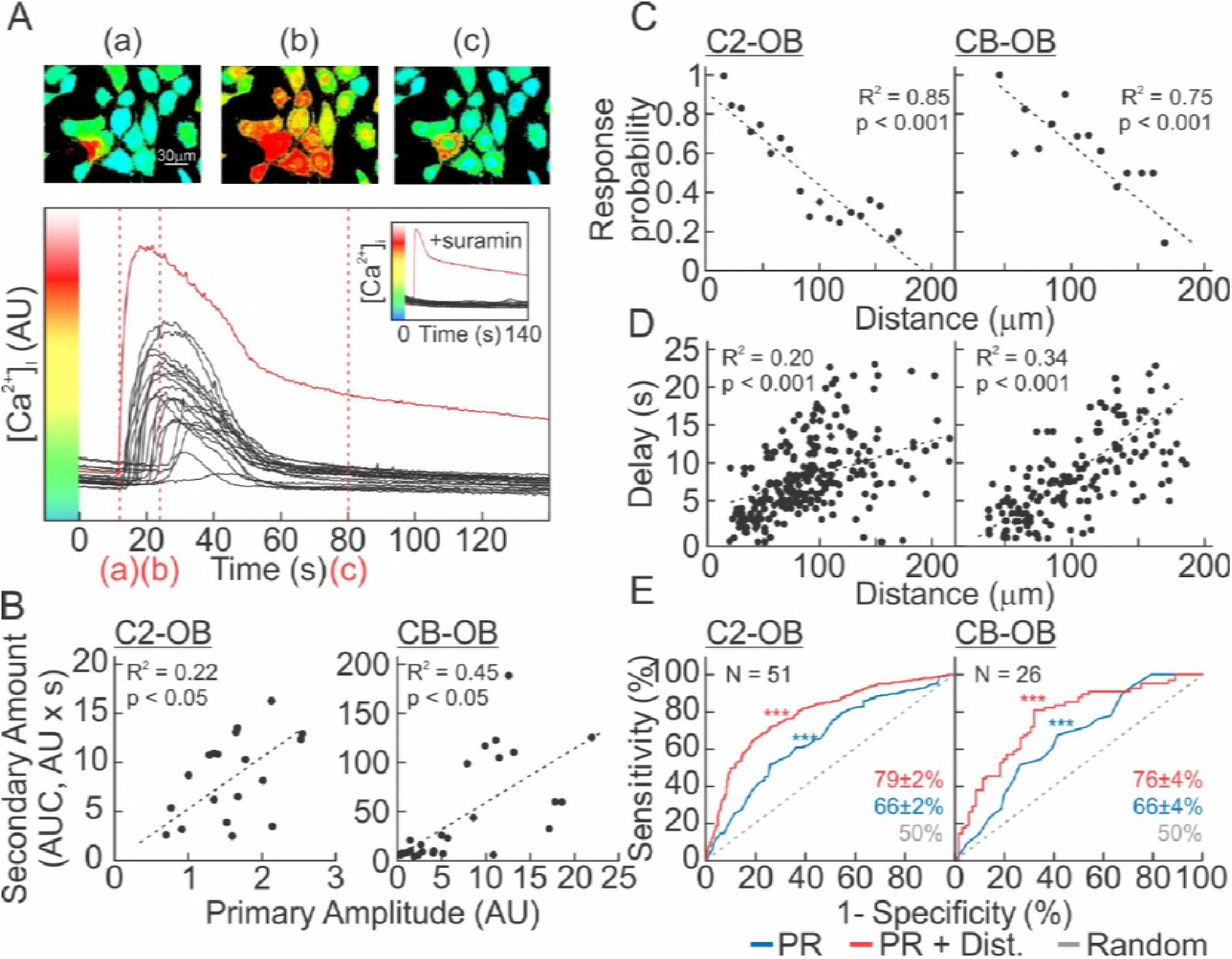
Mechanically-induced purinergic signaling conveys the magnitude and distance to stimulus. **(A)** Single Fura2-loaded C2-OB was mechanically stimulated by micropipette and [Ca^2+^]_i_ was recorded. *Top*: pseudocolored 340/380 Fura2 ratio images. *Bottom*: [Ca^2+^]_i_ recording of mechanically-stimulated (primary) cell (*red*) and neighboring (secondary) responders (*black*). *Red dashed lines*: Time points in top panel. *Inset*: Experiment performed in presence of P2 receptor antagonist suramin (100 μM). **(B)** Correlation between the primary response amplitude and area under curve (AUC) of secondary [Ca^2+^]_i_ elevations for C2-OB (*left*) and CB-OB (*right*). **(C-D)** Correlation between distance from source and response probability of neighboring cells **(C)** and delay to onset of response after mechanical stimulation **(D)** for C2-OB (*left*) and CB-OB (*right*). *Black dashed line*: linear regressions. **(E)** Receiver operating characteristic (ROC) curves demonstrate performance of logistic regression model in predicting incidence of secondary responses based on primary response (PR) parameters (*blue*), PR parameters and distance (dist., *red*) or chance alone (*grey*) for C2-OB (*left*) and CB-OB (*right*). Accuracies of logistic models ± SEM are shown, N indicates number of trials.

### 5.2 Mechanically-induced ATP release depends on the size and resealing kinetics of membrane disruption

We previously demonstrated that mechanical stimulation of an osteoblast generates non-lethal membrane injury, through which intracellular ATP is released (7). Membrane injury was then quickly repaired through PKC-dependent vesicular exocytosis, thus limiting ATP release (7). To quantify the size of membrane injury and the rate of membrane resealing, we first experimentally determined the characteristic half-time to repair *τ*_1/2_ using two independent methods. Fura2-loaded C2-OB cells were stimulated by a glass micropipette and the subsequent decline in Fura2 340 ex/510 em fluorescence were monitored until a plateau was achieved, thereby indicating membrane repair (**Fig 2A**, *top*). Additionally, quinacrine-loaded C2-OB cells were micropipette-stimulated and vesicular release was monitored as a proxy of the membrane repair process (**Fig 2A**, *bottom*). The repair time *τ*_1/2_, calculated as the exponential half-life, was 11 s (95% CI: 6 to 16) for Fura2-leakage assay (**Fig 2B**, *blue*) and 18 s (95% CI: 7 to 28) for vesicular release imaging (**Fig 2B**, *red*). We have also demonstrated using dye exclusion assays that the initial injury size is on the nanometer scale (7). We modelled the injury size *d* (*t*) as an initial membrane disruption *d*_0_ that closes with an exponential half-life *τ*_1/2_ (**Eq. 1**), and ATP release as diffusion through this injury (**Eq. 2-3**). At varying *d_o_* and *τ*_1/2_ (**Fig 2C**), ATP release was highest immediately after membrane disruption and declined exponentially as the size of injury decreased (**Fig 2D**). The total amount of ATP released ([*ATP*]_*total*_) was proportional to the initial injury size *d_o_* and inversely proportional to the characteristic repair time τ_1/2_ (**Fig 2E, F**). For a wide range of injury parameters, less than 30% of total cellular ATP was released and this was previously determined to be non-lethal to the cell (7) (**Fig 2E**). Unlike [*ATP*]_*total*_, the peak concentration of extracellular ATP ([*ATP*]_*peak*_) was solely determined by the initial injury size (**Fig 2G**), and ATP persistence at the source ([*ATP*]_*FWHM*_, duration that extracellular [ATP] was sustained above half-maximal [ATP]) was predominantly governed by the rate of membrane resealing (**Fig 2H**). Thus, the severity and dynamics of membrane repair differentially contribute to the peak, total amount and persistence of extracellular ATP concentrations.

**Figure 2.**
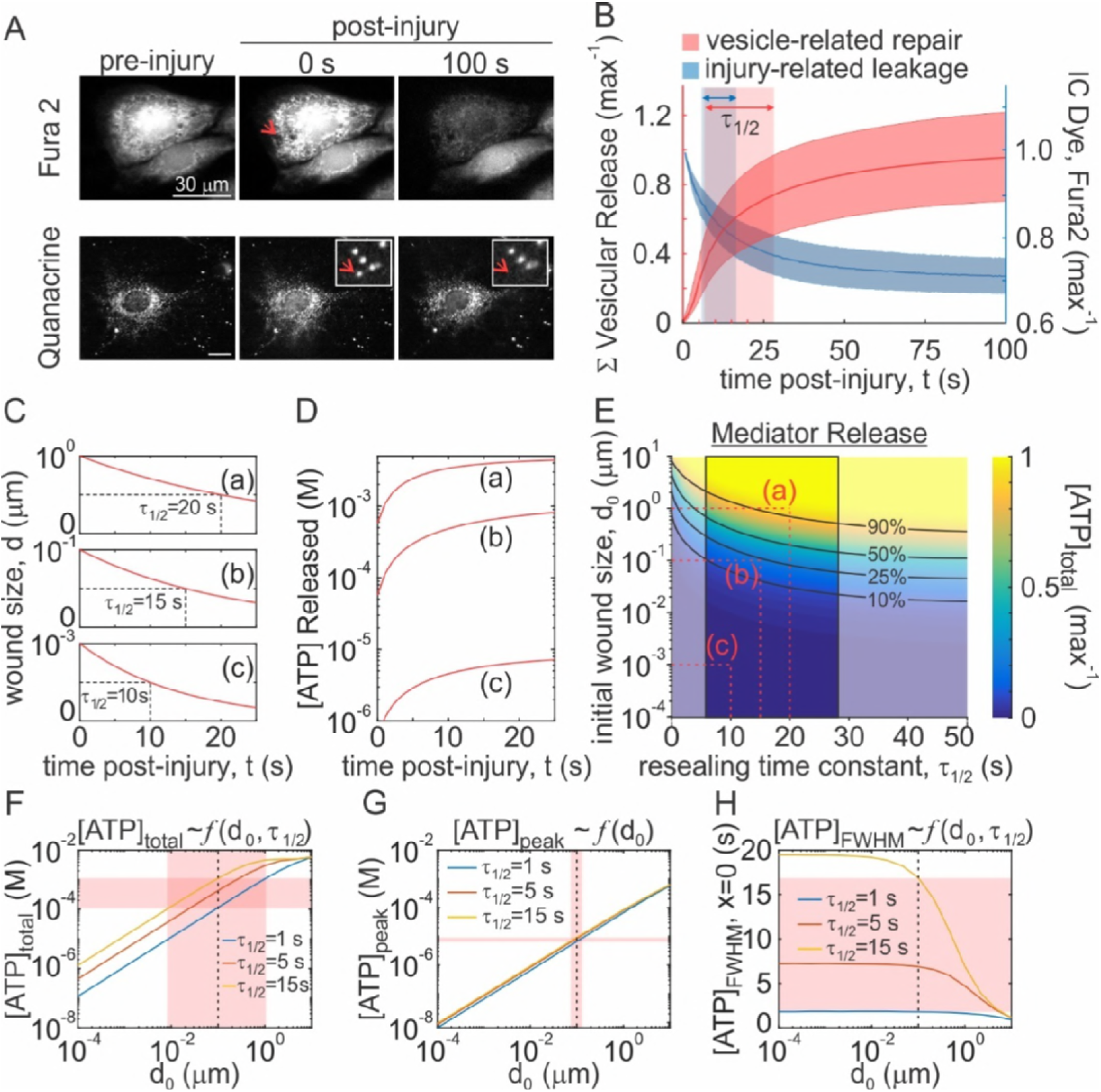
The size and resealing kinetics of the membrane injury determine the kinetics of ATP release. **(A, B)** Single Fura2- (**A**, *top*) or quinacrine- (**A**, *bottom*) loaded C2-OB was mechanically stimulated by micropipette and time-course of intracellular (IC) Fura2-leakage (**B**, *blue,* N=51) and cumulative vesicular release (**B**, red, N=15) was recorded. Data are means ± 95% confidence intervals (CI). *Shaded boxes*: 95% CI of time to half max (*τ*_1/2_) vesicular release (*red*) or Fura2-leakage (*blue*). **(C, D)** Simulated time-dependent changes in injury size *d*(*t*) **(C)** and ATP release **(D)** for indicated initial injury size *d*_0_ and the characteristic membrane repair time *τ*_1/2_. **(E)** Solution space for injury-related ATP release with respect to changes in initial injury size *d*_0_ and the characteristic membrane repair time *τ*_1/2_. *Black lines*: Isoclines for parameter pairs that yield equal percentage of total intracellular ATP release. *Red dashed lines*: Parameter pairs (a, b, c) used in **C, D**. *Outlined region*: Experimentally measured *τ*_1/2_. **(F-H)** Relationship between initial injury size *d*_0_ and total ATP released [*ATP*]_*total*_ **(F)**, peak extracellular ATP concentration[*ATP*]_*peak*_ **(G)** and full-width half max (FWHM) duration of extracellular ATP concentration [*ATP*]_*FWHM*_ at source **(H)** for *τ*_1/2_ = 1, 5, 15 s. *Red bands* show extent of influence *τ*_1/2_ has on ATP release parameters with respect to *d*_0_ = 100 nm (*dashed black line*).

### 5.3 Purinergic response probabilities in the presence of ATP and ADP

Mediator spillage through a membrane injury is nonspecific, therefore, any purines present in the cytosol will be released into the extracellular space in proportional amounts. We tested adenine purines ATP, ADP, AMP and adenosine for their ability to induce calcium responses (**Fig 3A**). Only ATP and ADP were capable of evoking [Ca^2+^]_i_ elevations in C2-OB cells, with ATP being an order of magnitude more potent than ADP (**Fig 3A, B**). When ATP and ADP were applied together in varying concentrations, response probabilities were dictated by the most potent nucleotide species present (**Fig 3C**). We modeled ATP and ADP diffusion profiles (**Fig 3D; Eq. 4, 5**) and related local ATP and ADP concentrations to response probabilities obtained from experimental data (**Fig. 3B, C; Eq. 7**), allowing us to generate a spatiotemporal map of secondary response probabilities (**Fig 3E**). Response probabilities for neighbouring cells were computed for a range of injury-related parameters (*d*_0_ = 10^−4^ to 10^1^ μm, *τ*_1/2_ = 1 to 20 s) and compared to experimental response probabilities obtained from single-cell micropipette stimulation recordings to identify the parameter space that was consistent with observations (**Fig S1A**). For characteristic repair times longer than 4 s, secondary response probabilities predicted for 50 nm to 1 μm injuries were consistent with observations (**Fig S1A**), with the best fit parameter set being *d*_0_ = 100 nm and *τ*_1/2_ = 15 s (**Fig 3F**). Thus, we demonstrated that P2 receptor-mediated response probabilities are governed by the most potent nucleotide present, and computationally estimated the injury parameters that recapitulate paracrine response probabilities that were measured *in vitro*.

**Figure 3.**
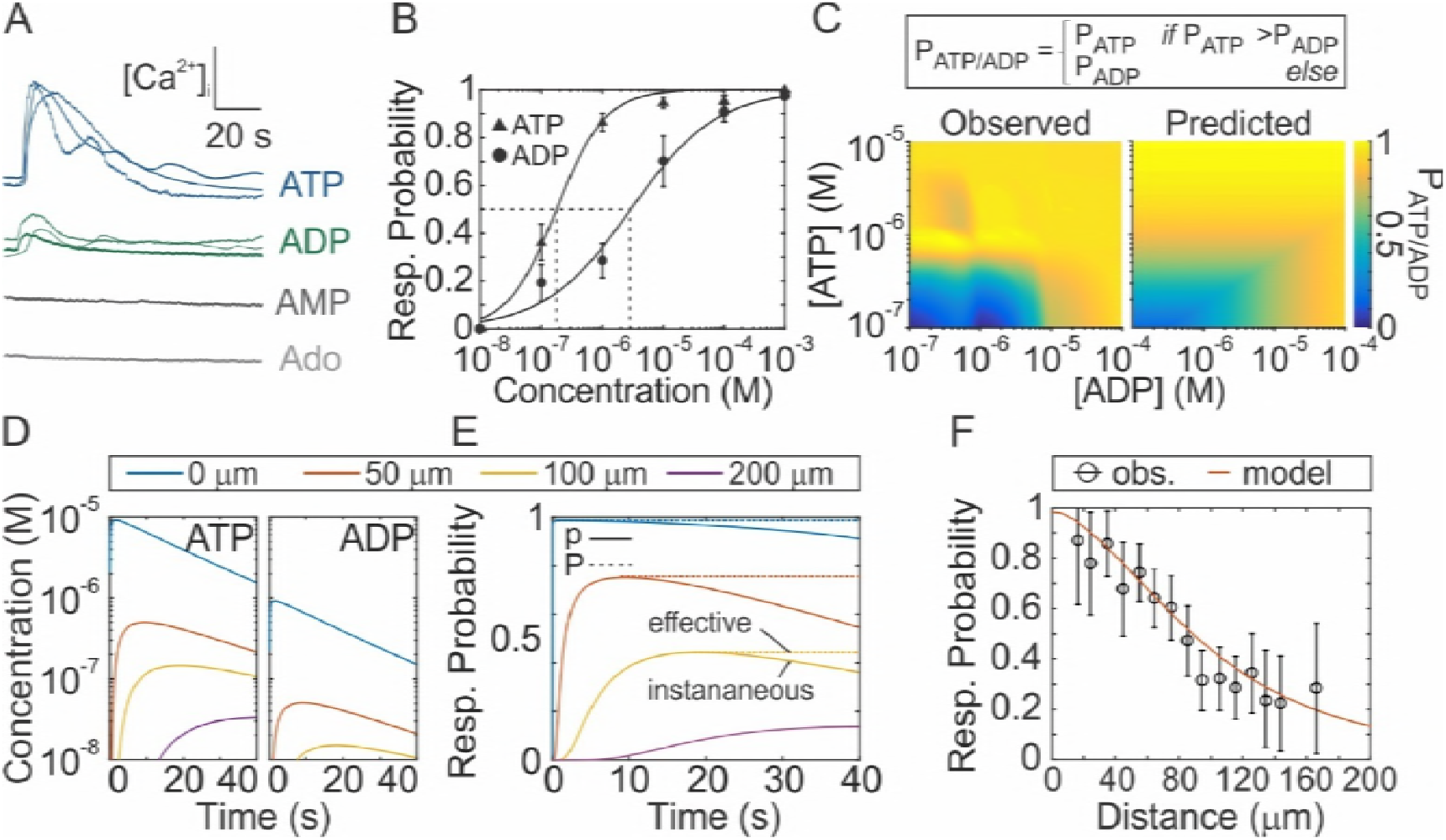
Dependence of secondary response probabilities on ATP and ADP release and diffusion. **(A-C)** Fura2-loaded C2-OB were stimulated by bath application of ATP, ADP, AMP or adenosine. **(A)** Representative [Ca^2+^]_i_ elevations stimulated by 1 μM ATP, 1 μM ADP, 10 μM AMP or 10 μM adenosine (Ado). **(B)** ATP and ADP dose-dependent response probabilities. Means ± SEM, N=6-8. *Fitted curves*: Hill functions (**Table S1**). **(C)** Response probabilities *P* of C2-OB stimulated with varying combinations of ATP and ADP. Empirical relationship between ATP/ADP and response probability (*top panel*) was established using observed data (*left surface plot*) and used to predict response probability (*right surface plot*). **(D-E)** ATP/ADP release and diffusion was numerically simulated (*d*_0_ = 100 nm, *τ*_1/2_ = 15 s, Eq. 4,5, **D**) and used to predict instantaneous (*p*) and effective (*P*) response probabilities (Eq. 7, **E**). **(F)** Comparison of predicted and observed distance-dependent response probabilities following mechanical stimulation of a single osteoblast at *x* = 0. Observed data are means ± SEM, N=51.

### 5.4 Purinergic response times in the presence of ATP and ADP

We next examined whether diffusion of ATP and ADP could account for the delay between mechanical stimulation and subsequent paracrine responses. ATP and ADP diffusion coefficients were reported to be ~347 and 377 μm^2^s^−1^, respectively (24-26). We determined the apparent diffusion coefficient *D_α_* (**Fig 4A**) from the slope of the linear relationship between secondary response delay times and squared distances from the source as 222 μm^2^s^−1^ (95% CI: 214 to 239), which was significantly lower than expected. To examine whether the discrepancy between apparent and expected diffusion coefficients was related to the presence of different nucleotides, we performed mechanical stimulation of the primary cell and monitored the time to response in neighboring cells in the presence of pyruvate kinase and phosphate-donor phosphoenolpyruvate, which enzymatically converted extracellular ADP to ATP, or in the presence of hexokinase which converted ATP to ADP (**Fig 4B**). Pyruvate kinase had no effect, while hexokinase further reduced the apparent diffusion coefficient (**Fig 4C, D**), contrary to what was expected based on the lower molecular weight of ADP compared to ATP. We hypothesized, that in addition to diffusion, the reaction times (*t_rxn_*) of ATP- and ADP-mediated responses may exhibit purine-species dependent differences. Applying ATP, ADP or their combination to Fura2-loaded C2-OBs, we estimated the average time delay between purine application (approximated as time of first discernable cell response) and subsequent cell responses (**Fig 4E**). ADP-mediated reaction times were consistently slower than ATP-mediated reaction times (**Fig 4F**). Importantly, when ATP and ADP were applied together in varying concentrations, the most potent nucleotide governed the observed reaction time (**Fig 4G**). To account for the influence of P2 response times, we formulated the problem as a response time model (21). Using numerically simulated ATP and ADP concentration profiles to determine local ATP and ADP concentrations (**Eq. 4, 5**), and experimental dose-dependencies of *t_rxn_* (**Fig 4F, G**), we derived instantaneous response rates *λ*(*x*, *t*) as the reciprocal of *t_rxn_* (**Fig 4H; Eq. 8)**. This allowed us to define the time- and distance-dependent proportion of non-responding cells (**Fig 4I; Eq. 9, 10**), and to predict secondary response times *T* at different distances from the primary cell (**Fig 4J; Eq. 11**). To identify the injury-related parameter space (*d*_0_, *τ*_1/2_) that was consistent with observations, the combined bias between observed and predicted reaction times and response probabilities was determined for *d*_0_ = 10^−4^ to 10^1^ μm and *τ*_1/2_ = 1 to 20 s (**Fig S1)**. We showed that *τ*_1/2_ > 4 s and *d*_0_ = 100 to 500 nm generated predictions consistent with observed outcomes (**Fig S1)**, with *d*_0_ = 100 nm and *τ*_1/2_ = 15 s remaining the best fit parameter set (**Fig 4J-K**). Moreover, the predicted spatiotemporal probability distribution of responses *f*(*x*, *t*) adequately recapitulated the variation in observed paracrine response times (**Fig 4K; Eq. 12**). Thus, accounting for ATP and ADP diffusion and reaction times is sufficient to quantitatively describe the spatiotemporal pattern of secondary purinergic responses.

**Figure 4.**
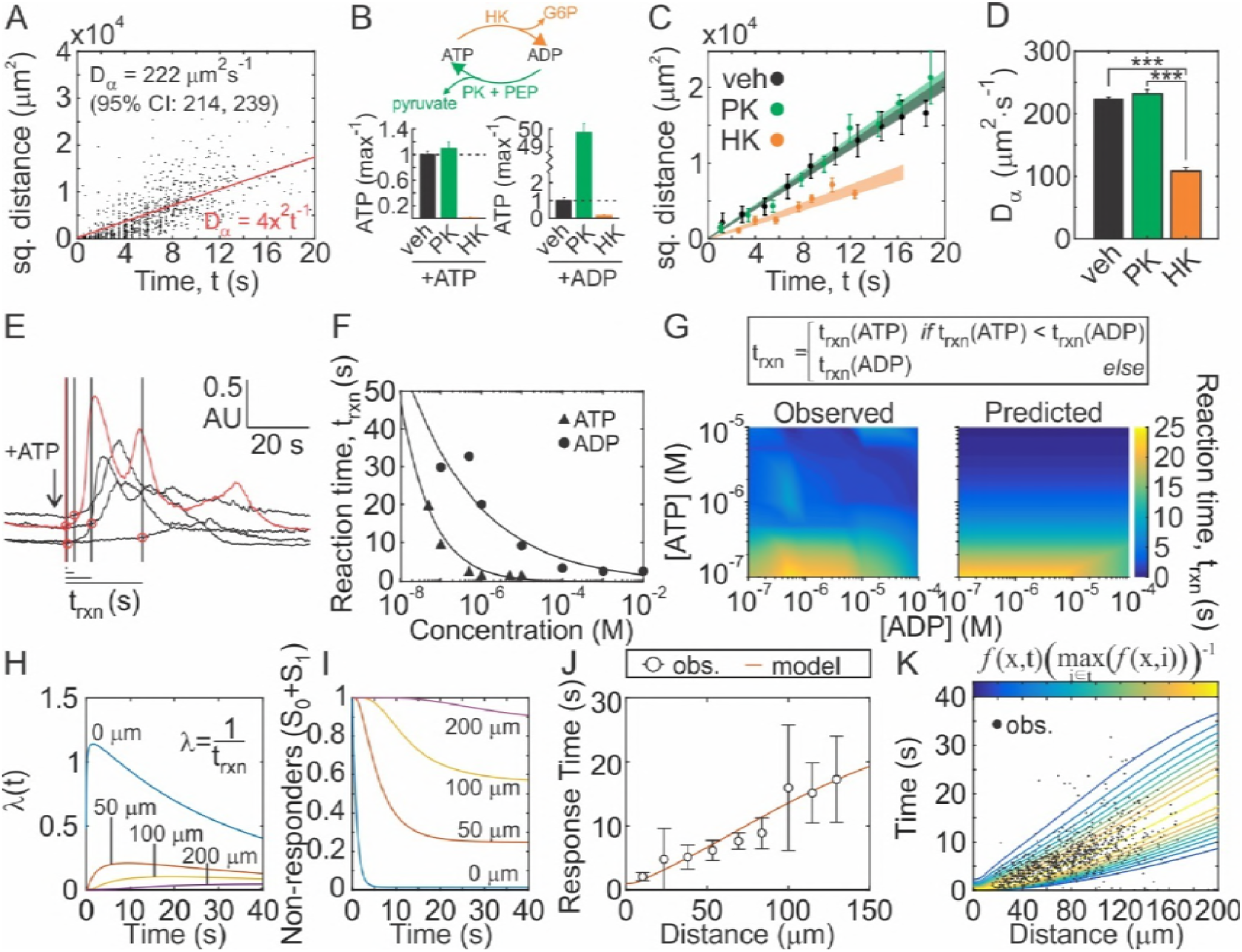
Response-time modeling predicts spatial and temporal distribution of paracrine responses. **(A)** Relationship between measured secondary response times and squared distance from source following micropipette stimulation of single Fura2-loaded C2-OB. *Red line*: Apparent diffusion coefficient D_α_ estimated by linear regression. **(B)** 1 μM ATP (*left*) or ADP (*right*) solutions were incubated with vehicle, pyruvate kinase (PK) + phosphoenolpyruvate (PEP) or hexokinase (HK) for 5 min and [ATP] was measured. Means ± SEM, N=3. **(C, D)** Fura2-loaded C2-OB was mechanically-stimulated by micropipette in presence of vehicle, PK (+PEP) or HK and relationships between secondary response times and squared distance from the source were evaluated **(C,** *shaded bands*: 95% CI). D_α_ were determined by linear regression **(D)**. Means ± SEM, ***p<0.001 assessed by ANOVA and Bonferroni test, N=10-15. **(E-G)** Fura2-loaded C2-OB were stimulated by bath application of ATP, ADP, or their combination. **(E)**Representative [Ca^2+^]_i_ traces show reaction times *t_rxn_* for ATP-stimulated responses (1 μM, *black arrow*). *Red trace*: first detected response, *black traces*: subsequent responses*, black lines/red circles*: response times. **(F)** ATP/ADP dose-dependences for *t_rxn_*. *Black lines*: fitted exponential functions (**Table S1**). **(G)** Empirical relationship between ATP/ADP and *t_rxn_* (*top panel*) was established using observed data (*left surface plot*) and used to predict *t_rxn_* (*right surface plot*). **(H, I)** ATP/ADP release and diffusion was numerically simulated (*d*_0_ = 100 nm, *τ*_1/2_ = 15 s, Eq. 4,5) and used to predict instantaneous reaction rates *λ* (Eq. 8, **H**), space- and time-dependent proportion of non-responding neighboring cells *S* = *S*_0_ + *S*_1_ (**I**). **(J, K)** Comparison of predicted and observed secondary response times **(J)** and spatiotemporal response probabilities represented as probability density function *f*(*t*) (Eq. 12) overlaid with observed data **(K)**. Observed (obs.) data are means ± SEM in **J** and raw data in **K**, N=51.

### 5.5 Severity of membrane injury and repair dynamics are differentially decoded by secondary purinergic responses

To characterize the relationship between the mechanical stimulus and the involvement of neighbouring responders, we introduced parameters describing the degree of response (recruitment parameters) and the temporal characteristics of the response (timing parameters). Response recruitment parameters described the extent of involvement from the neighbouring population and included the signal radius *R_signal_*; distance at which the paracrine response probability declined below 5% (**Fig 5A**, *left*), and the total response *R_AUC_*; area under the distance-dependent response probability curve (**Fig 5A**, *right*). Timing parameters described the temporal characteristics of the paracrine responses and included the signal velocity; *V_signal_* average speed at which the paracrine signal propagated (**Fig 5B**, *left*), and temporal synchrony *t_sync_*; full-width half-max (FWHM) of the response time probability distribution *f* (**Eq. 12**) at a given distance (**Fig 5B**, *right*). We found that response recruitment parameters were governed by the total amount of ATP released regardless of whether ATP was spilled from a severe injury (large *d*_0_) or from a slowly repaired minor injury (small *d*_0_, **Fig 5C**). In contrast, response velocity and synchrony were predominantly governed by the peak concentration of ATP achieved following injury, with a more pronounced dependence at higher peak ATP concentrations due to more severe injuries (**Fig 5D**). These data suggest that spatiotemporal variations in ATP concentration profiles differentially encode response recruitment and timing parameters, such that peak ATP concentrations govern response timing and the total amount of ATP released governs the degree of involvement of the neighbouring population. Thus, we postulate that neighboring cell populations can distinguish between severe and minor mechanically-induced cell injuries that release similar amount of ATP (due to diverse repair dynamics).

**Figure 5.**
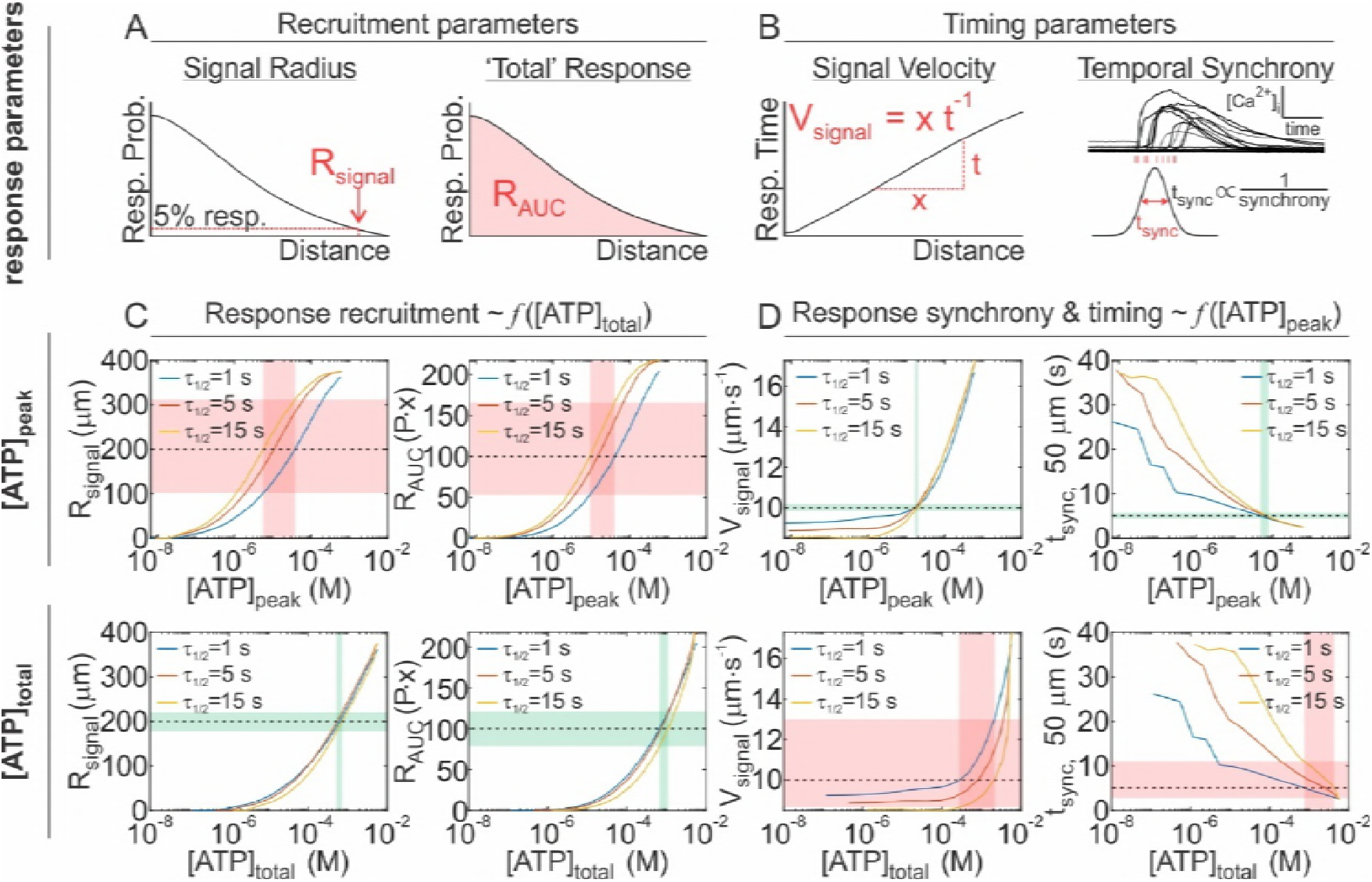
Relationship between ATP release and recruitment and timing of secondary [Ca^2+^]_i_ responses. **(A, B)** Schematics of response recruitment parameters: signalling radius (*R_signal_*, **A**, *left*) and total response (*R_AUC_*, **A**, *right*), and response timing parameters: signal velocity (*V_signal_*, **B**, *left*) and temporal synchrony (*t_sync_*, **B**, *right*). **(C, D)** Relationship between ATP release parameters [*ATP*]_*peak*_ (*top*) or [*ATP*]_*total*_ (*bottom*) and response recruitment parameters *R_signal_* (**C**, *left*) and *R_AUC_* (**C**, *right*) or response timing parameters *V_signal_* (**D**, *left*) and *t_sync_* (**D**, *right*) for *τ*_1/2_ = 1, 5, 15 s. *Dashed black lines*: Reference for *R_signal_* = 200 μm (**C**, *left*), *R_AUC_* = 100 P·x (**C**, *right*), *V_signal_* = 5 μm s^−1^ (D, *left*) and *t_sync_* = 5 s (**D**, *right*). *Shaded bands*: Show extent of influence of *τ*_1/2_ (*red*: large influence, *green*: minimal influence) on response parameters at dashed-line references.

### 5.6 Characterizing extracellular degradation of ATP released by a mechanically-stimulated osteoblast

In the extracellular space there are ecto-nucleotidases that hydrolyse ATP and ADP to various metabolites. In the context of bone, there are several resident cell types, including osteoblasts, osteoclasts and bone marrow cells, which can potentially contribute to extracellular ATP degradation. We found that osteoblast-(C2-OB) and osteoclast precursors-like (Raw 264.7) cells metabolized extracellular ATP similarly with rate constants of 0.049 min^−1^ (95% CI: 0.043 to 0.055) and 0.048 min^−1^ (95% CI: 0.044 to 0.042), respectively, while erythrocyte-like (K562) cells hydrolyzed ATP significantly faster with a rate constant of 0.096 min^−1^ (95% CI: 0.085 to 0.11) (**Fig 6A, Table S1**). The degradation rate was independent of cell density (**Fig S2**). In C2-OB cells, ATP degradation was partially inhibited by ecto-nucleotidase inhibitor ARL 67156 (**Fig 6B**, *blue*), however, it did not influence the responding fraction of neighbouring cells following mechanical stimulation of an individual Fura2-loaded C2-OB (**Fig 6C**). We also found that ATP degradation was potently inhibited by alkaline phosphatase inhibitor orthovanadate (**Fig 6B**, *red*); however, it altered ATP-mediated responses independently of its effects on degradation rendering signal propagation experiments in the presence of orthovanadate uninterpretable (**Fig S3**). The presence of ADP-to-ATP converting enzyme pyruvate kinase did not affect paracrine responses (**Fig 6D, E**, *green*). To examine the potential effect of fast ATP degradation, we used ATP-to-ADP converting enzyme hexokinase, which resulted in a significant reduction in the response radius and fraction (**Fig 6D, E**, *orange*). Mechanically-induced ATP and ADP concentrations profiles were also computationally simulated in the presence of extracellular nucleotide degradation (*k_ATP_* = 0.05 min^−1^, *k_ADP_* = 0.25 *k_ATP_*, **Eq. 4-5**) and paracrine response probabilities were determined (**Fig 6F**). Consistent with experimental findings, numerical simulations demonstrated that endogenous levels of nucleotide degradation are too slow to influence purinergic signals released by a single mechanically-injured cell.

**Figure 6.**
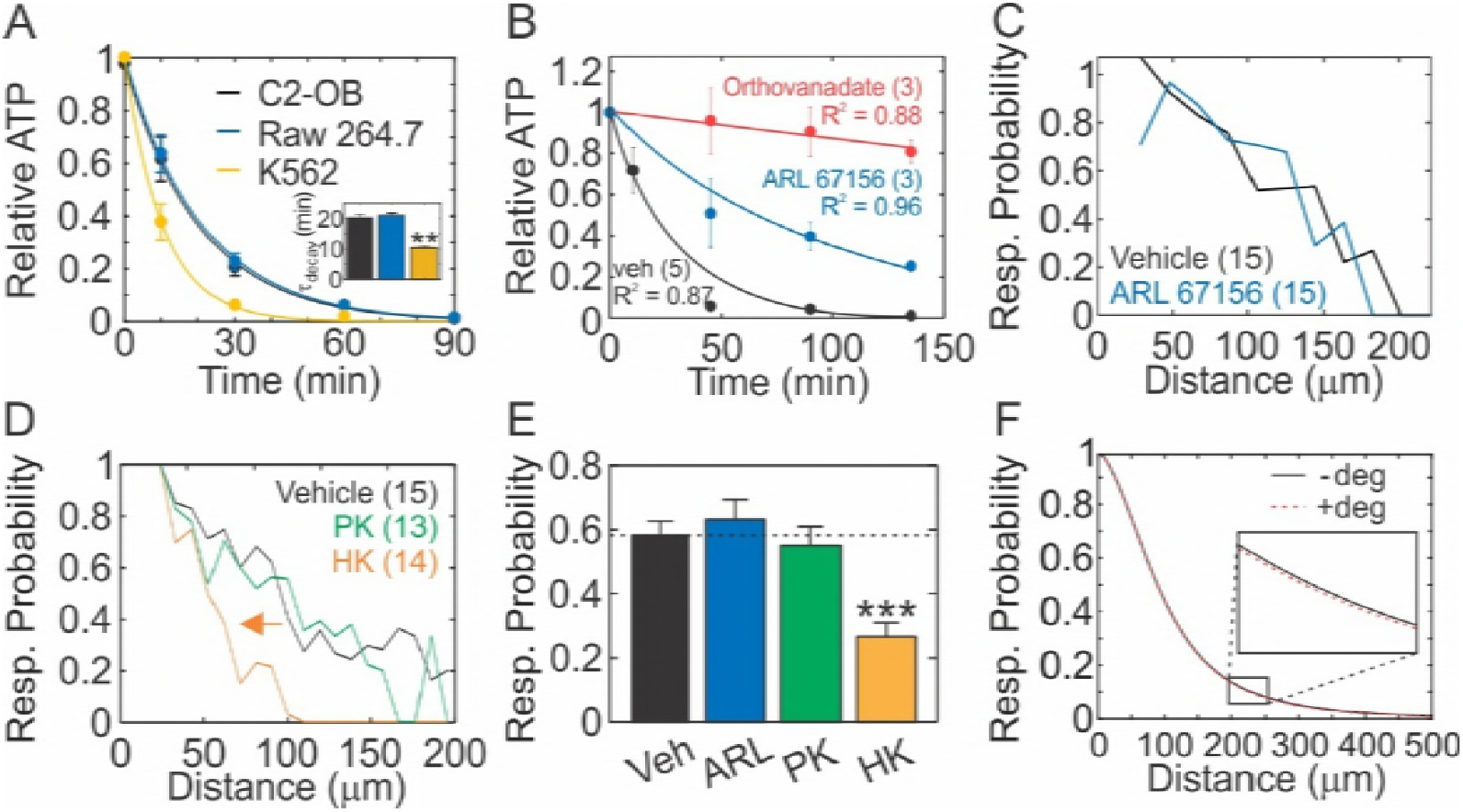
Influence of ATP degradation on mechanotransductive paracrine responses following single-cell injury. **(A)** 1 μM ATP was added to cultures of C2-OB osteoblasts, Raw 264.7 osteoclast-precursors and K562 erythrocyte-like cells and ATP degradation was measured. *Inset*: decay times constants *τ_decay_*. **(B)** Degradation of 1 μM ATP by C2-OB in the presence of ecto-nucleotidase inhibitor ARL 67156 (10 μM) or alkaline-phosphatase inhibitor orthovanadate (10 μM). For A, B, data are means ± SEM, *solid curves*: fitted exponential functions (**Table S1**). **(C-E)** Paracrine response probabilities following micropipette stimulation of single Fura2-loaded C2-OB cell in the presence of 10 μM ARL 67156 (ARL) **(C)**, 20 U/mL pyruvate kinase + excess phosphoenolpyruvate (PK) or 20 U/mL hexokinase (HK) **(D)** were quantified **(E)**. Sample size N in parentheses. Means ± SEM, ***p<0.001 assessed by ANOVA and Bonferroni test. **(F)** Numerically simulated paracrine response probabilities in the presence or absence of extracellular ATP degradation using degradation rate constants k_ATP_ measured in C2-OB cells. *Simulation parameters*: *k_ATP_* = 0 (degradation absent, - deg) or 0.05 min^−1^ (degradation present, + deg), *d*_0_ = 100 nm, *τ*_1/2_ = 15 s.

### 5.7 Propagation of purinergic signal resulting from a tissue-level injury

Physiological mechanical stimuli affect multiple cells occupying a tissue area of a certain size and geometry. This results in the release of purinergic mediators from numerous sources in the tissue, unlike the single cell experimental setup that we have considered thus far. To investigate how the extent and geometry of an injury affects paracrine signal propagation, we considered three injury geometries: (i) point source, consisting of one or 40 injured cells positioned at the center of the signalling axis (**Fig 7A**, *left*), (ii) linear source, consisting of 40 or 1600 injured cells evenly distributed along a 2000 μm linear plane perpendicular to the signalling axis (**Fig 7A**, *middle*), and (iii) half-field source consisting of 800 or 8000 cells evenly arranged as a series of point sources over a 2000 × 4000 μm^2^ surface area with the signalling axis running along the 2000 μm axis of injured cells and an additional 2000 μm distance with no injured cells (**Fig 7A**, *right*). For each of these tissue-level injury geometries, we assumed simultaneous and identical cell-level injury (*d*_0_ = 100 nm, *τ*_1/2_ = 15 s) and examined paracrine response probabilities in the presence and absence of extracellular nucleotide degradation (*k_ATP_* = 0.05 min^−1^, *k_ADP_* = 0.25 *k_ATP_*, **Fig 7B**). Since certain P2 receptors (e.g., P2Y1, P2Y12 and P2Y13) are predominantly sensitive to ADP (28), we also isolated the ADP-mediated component of the response probability (**Fig 7C**). The purinergic response was consistently highest at the site of injury and declined at increasing distances from the injury (**Fig 7B, C**). For all injury geometries, the more cells were initially injured, the more neighboring cells exhibited P2 responses (**Fig 7B, C**). The effect of extracellular nucleotide degradation became apparent for larger tissue-level injuries, especially when higher ATP concentrations (due to more cell injury) were sustained for longer (due to dispersed injury geometry). In these cases, ATP degradation served to attenuate total paracrine responsiveness while increasing the ADP-mediated signalling component (**Fig 7B, C**, *red vs. black curves*). Importantly, the effects of extracellular nucleotide degradation were not observed at the site of injury, but rather at increasing distances from the source. Thus, scaling a single-cell injury to a tissue-level injury with varying geometries revealed that ATP- and ADP-mediated components of the purinergic response reflect the extent of injury and differentially change with increasing distance from the injury site.

**Figure 7.**
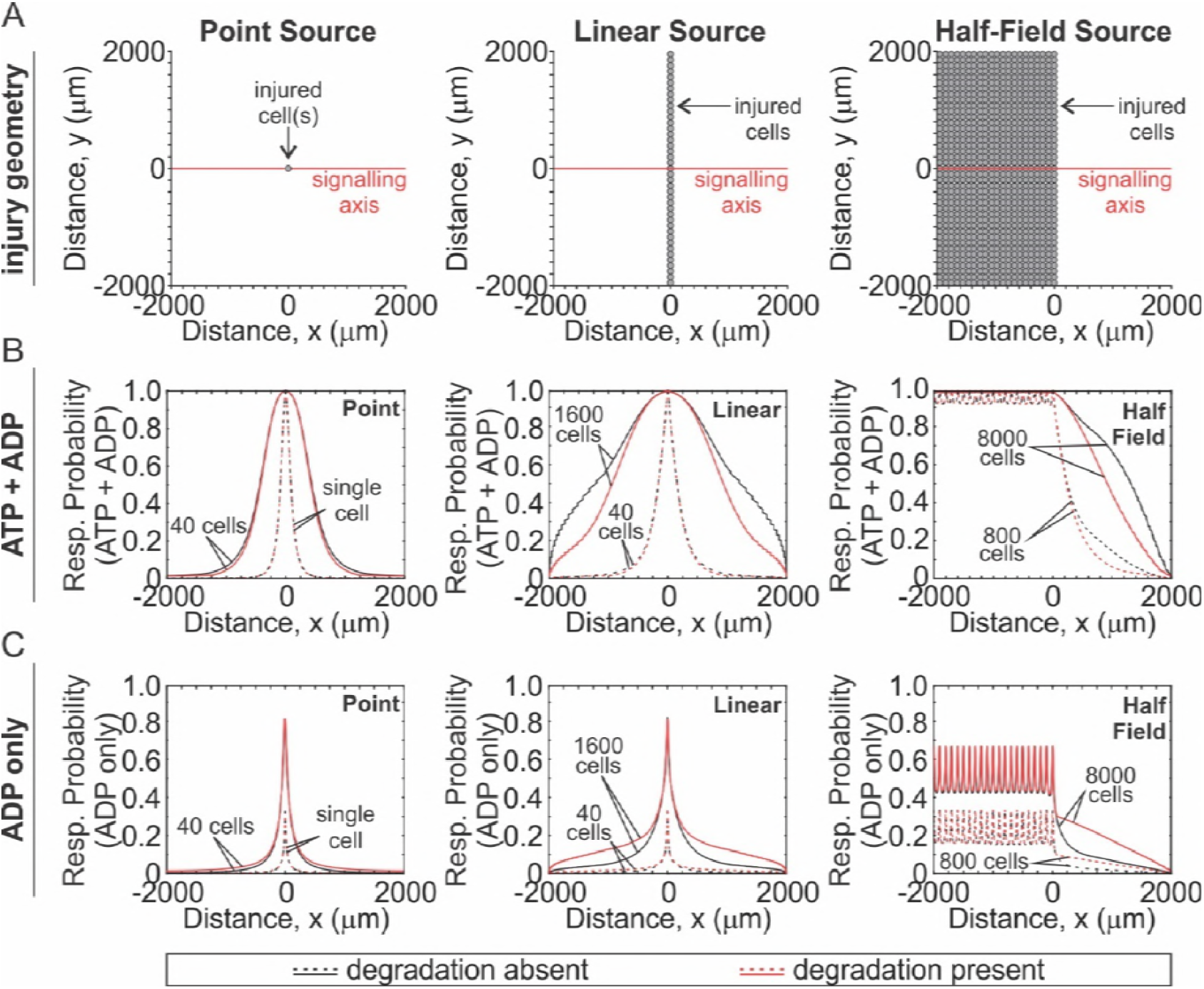
Contribution of ATP degradation and ADP to mechanotransductive paracrine signaling following tissue-level injury. ATP and ADP release and diffusion from point- (*left*), linear- (*middle*) and half-field- (*right*) source injuries were numerically simulated (Eq. 4,5, **A**) and contributions of ATP and ADP **(B)** or ADP alone **(C)** to paracrine response probabilities were determined for varying number of injured cells in the presence (*red*) or absence (*black*) of extracellular ATP and ADP degradation. *Simulation parameters*: *k_ATP_* = 0 (degradation absent) or 0.05 min^−1^ (degradation present), *k_ATP_* = 0.25 *k_ATP_*, *d*_0_ = 100 nm, *τ*_1/2_ = 15 s.

## 6 Discussion

### Overview

The goals of this study were (i) to determine how mechanically-induced cell membrane injury and repair govern ATP and ADP release dynamics, (ii) to characterize ATP and ADP spatiotemporal diffusion profiles and (iii) to understand the basic principles dictating the timing and responsiveness of neighboring (non-stimulated) cells to the presented levels of ATP and ADP. For each elementary step (ATP/ADP release, diffusion and degradation, and P2 responses in neighboring cells), we developed mathematical descriptions and validated each experimentally. Finally, using the combined model of ATP-driven mechanotransduction, we scaled a single-cell injury to a tissue-level injury with varying geometries and described how neighboring cell populations detect the magnitude of injury and discern their position relative to the injury site. Thus, we were able to identify specific features of the ATP/ADP spatiotemporal signal that encode the severity of mechanical stimulus, and distance to stimulus, as well as the state of tissue mechano-resilience.

### ATP release due to mechanical injury

Several groups have demonstrated that mechanically-induced membrane disruptions occur routinely under physiological conditions in different tissues, including muscle fibers (29), gastrointestinal tract (30), heart (31), aorta (32) and bone (7, 8). Despite these injuries, cell death is minimal because cells can rapidly repair these injuries (33). Membrane disruption results in nonspecific spillage of intracellular contents, which includes signalling mediators, such as growth factors (31, 34) and ATP (7, 35, 36). The extent of this spillage is determined by (i) membrane integrity (described by parameter *d*_0_) and (ii) the rate of membrane repair which actively limits the amount of spillage (parameter *τ*_1/2_). Importantly, both membrane integrity and repair rates are sensitive to pharmacological interventions along the Ca^2+^/PLC/PKC axis (7, 37) and are potentiated by prior exposures to mechanical stimulation (7, 38, 39). We described the dynamics of membrane injury and repair using an empirically-validated exponential function and demonstrated that injuries with an initial size of *d*_0_~100 nm that resealed with a half-time of *τ*_1/2_~15 s were consistent with experimentally observations. Moreover, predicted ATP concentrations at the surface of the mechanically-stimulated cell (~10 μM) were consistent with experimental measurements (0.05-80.5 μM) (7). Modelling ATP release through a localized injury was based on the theoretical description developed for transport process of fluorescent dyes though membrane pores (9). We showed that the total amount of ATP released depended on both the severity of mechanically-induced injury *d*_0_ and rate of membrane repair *τ*_1/2_, while the peak extracellular ATP concentration was determined by the severity of the injury alone. Since the P2 receptor network consists of 15 receptor subtypes with ATP affinities covering over a million-fold range of ATP concentrations (16, 17), the peak ATP release will determine the degree of activation of low-affinity P2 receptors [e.g. P2X7; ec50 = 1.9 mM (human), ec50 = 130 μM (rat) (16)]. In contrast, the total amount of ATP released will determine the area over which high-affinity P2 receptors are activated [e.g. P2Y2; ec50 = 200 nM (human), ec50 = 2.9 μM (rat) (16)]. Thus, the P2 receptors network can decipher the information about the extent of the injury and the rate of repair encoded in the dynamics of ATP release. Importantly, these mechanistic insights are applicable to any form of stimulus-related mediator release which can be described in terms of its magnitude and temporal dynamics, including conductive and vesicular ATP release.

### Signal propagation through ATP diffusion and degradation and P2 responses in neighboring cells

Following release into the extracellular space, ATP diffuses and is concurrently metabolized by ecto-nucleotidases (40). ATP metabolites include ADP, AMP and adenosine, which may also stimulate calcium responses (14, 41-43). Moreover, since mediator spillage through a membrane injury is nonspecific, purines present in the cytosol will be released together with ATP in proportional amounts. Of all the tested purine metabolites, ADP was identified as potential mediator capable of stimulated calcium responses in C2-OB, consistent with prior work (44, 45). ATP degradation half-time in the extracellular space was 20 min (95% CI: 17 to 25) for C2-OB cells, compared to ~10 min by primary rat osteoblasts (46), and ~5 min by primary murine osteoblasts (47). However, in osteoblasts ATP degradation was too slow to affect purinergic transmission initiated by a single cell, contrary to other cell types in which paracrine responses following single cell mechanical stimulation were potentiated when ecto-nucleotidases were inhibited, such as in bovine endothelial cells treated with ARL-67156 (48, 49), or in human keratinocytes with siRNA-silenced NTPDase2 (50). Nevertheless, since most ecto-nucleotidases have a micromolar affinity for ATP (40), low hydrolysis rates were expected for the amounts of ATP released by a single mechanically stimulated osteoblast. Spatiotemporal ATP/ADP concentration profiles following mechanical stimulation were computationally simulated by coupling injury-related ATP release to radial diffusion. We did not use direct ATP measurements to validate the mathematical model of diffusion and degradation due to limited sensitivity and spatial resolution. Instead, we validated the model using [Ca^2+^]_i_ recordings of secondary responders following micropipette stimulation of a single osteoblast. Surprisingly, we have found that the apparent ATP diffusion coefficient derived from these [Ca^2+^]_i_ recordings was lower than previously reported (24-26). To account for this discrepancy, we built upon a response-time modeling framework to predict the probability and timing of secondary [Ca^2+^]_i_ elevations (21). Not only did response-time modeling account for the high degree of variability observed in intercellular [Ca^2+^]_i_ response times, but it allowed us to predict experimentally observed propagation velocities of ~10 μm/s, consistent with those reported by others (51). Accounting for response times resolved the discrepancy between observed and expected ATP diffusion coefficients that was shown by us and others (52). Importantly, we demonstrated that finite wave propagation was sufficient to explain observed intercellular [Ca^2+^]_i_ elevations in osteoblasts, without need for regenerative propagation terms (e.g., ATP-mediated ATP release) that were proposed in airway epithelia (53) or astrocytes (54). Thus, we have demonstrated that ATP/ADP diffusion and degradation as well P2 receptor response times collectively explain the timing of purinergic mechanotransductive events in osteoblasts.

### Mechanical information encoding

Throughout this study, we investigated which mechanotransductive features can communicate mechanically-relevant information to non-stimulated neighboring cells. We identified three such features that conveyed differential information about the mechanical stimulus: (i) the total amount of ATP released, (ii) the peak extracellular [ATP] and (iii) ADP-mediated signaling component. The total amount of ATP released, regulated by the injury severity (*d*_0_) and repair dynamics (*τ*_1/2_), determined the P2 receptor signaling radius and the overall number of responders, thus governing the extent to which the neighboring population was recruited. The second feature was the peak extracellular concentration of ATP which reflected the severity the individual cell injury (*d*_0_ only) and strongly influenced signal propagation velocity and temporal synchrony. Peak [ATP] allowed cells to discriminate between severe (large *d*_0_) and minor (small *d*_0_) injuries that led to the release of similar total amounts of ATP due to differences in injury repair dynamics (e.g. fast repair for large injury, slow repair for small injury). Therefore, severe injuries were associated with higher peak ATP concentrations that were capable of recruiting low-affinity P2 receptors (e.g. P2X7), and also synchronicity of responses at the population-level (synchronous for large injuries, asynchronous for small injuries), which is important for the coordination of tissue-level behaviors, such as arterial contraction and vasomotion (55), epithelial wound closure (56) and osteocytic network communication during *in vivo* mechanical loading (57). The third signaling feature that we identified was ADP-mediated signaling which became pronounced only when tissue-level injuries were considered. In the case of a single cell injury, the amounts of ADP released and produced from ATP degradation were too low to induce paracrine responses. However, this was no longer the case when the extent of injury was increased to include multiple cells, resulting in more ADP release and production by ATP degradation. The higher rate of ATP degradation following tissue-level injury was associated with overall dampening of paracrine responses, especially at further distances from the injury site, and the increase in ADP-mediated signaling was proportional to the extent of the injury. The observed injury-related increase in ADP-mediated signalling is supported by prior work in which ADP-sensitive P2Y1 (14) and P2Y13 (47) receptors were implicated in mechanotransductive signalling in osteoblasts. Thus, our findings support a model in which mechanical information is encoded within ATP/ADP diffusion waves are differentially decoded by low affinity P2 receptors, high affinity P2 receptors and ADP-sensitive P2 receptors (**Fig 8**).

**Figure 8.**
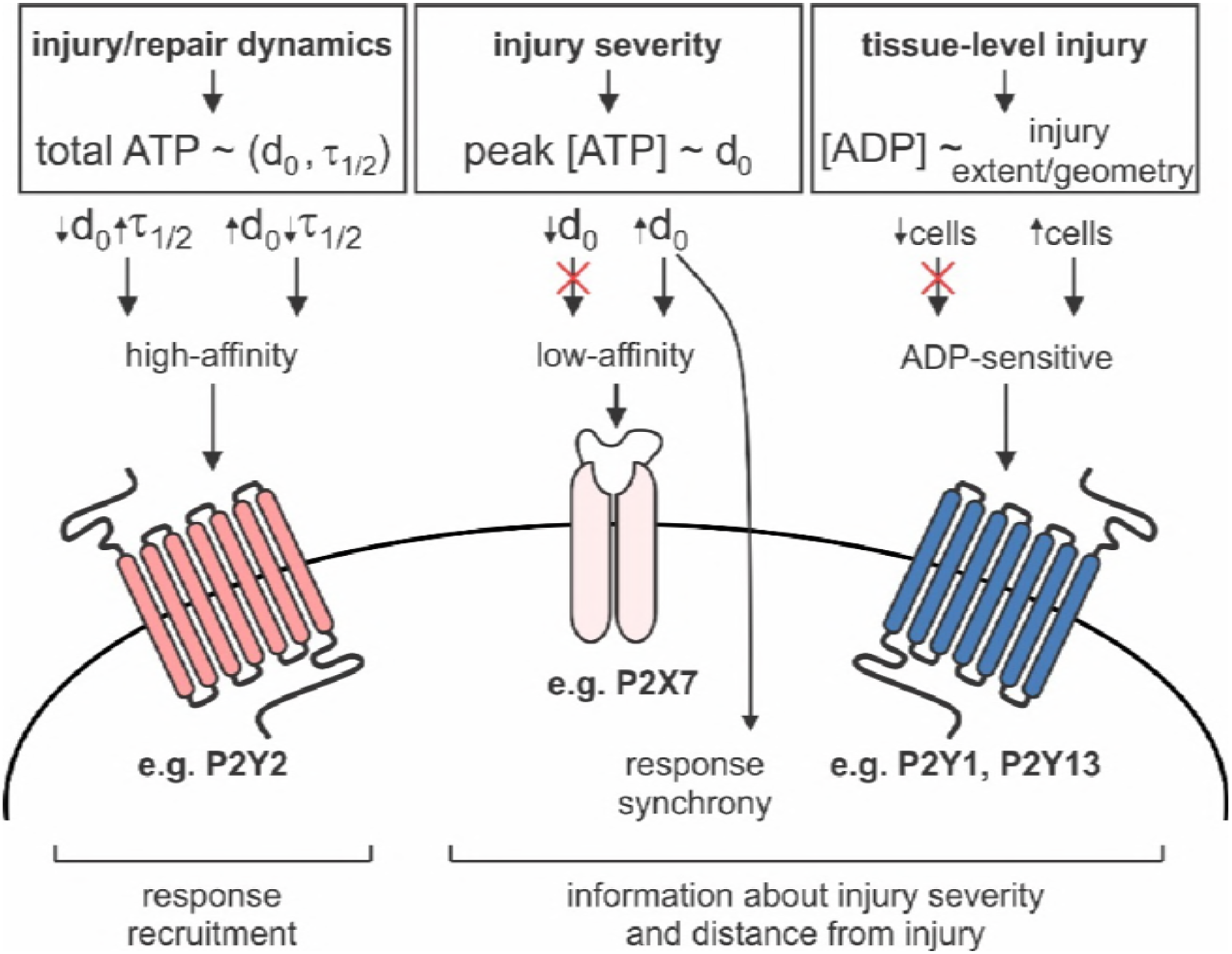
Proposed model of mechanical information decoding by the P2 receptor network. *Left*: Single cell membrane injury (*d*_0_) and repair (*τ*_1/2_) dynamics regulate the total amount of ATP released and the extent of recruitment of neighboring cells through high-affinity P2 receptors. *Middle*: Severity of injury (*d*_0_) is conveyed to neighboring cells through peak extracellular ATP concentrations which influence low-affinity P2 receptor signaling and timing of responses. *Right*: Extent and geometry of tissue-level injury determines amount of ADP released and produced by ATP degradation, and thus stimulation of ADP-sensitive P2 receptors.

### Concluding remarks

In the current study, we developed and validated a theoretical model to investigate how extracellular ATP and ADP released by mechanical stimulation encode information about the position and severity of the mechanical stimulus, and how this information can be subsequently decoded at the level of the paracrine responses. Our findings demonstrate that mechanotransductive purinergic signalling fields are tuned by the severity of injury and dynamics of repair, thereby enabling neighbouring cellular populations to evoke position- and injury-appropriate responses.

## Author Contributions

Study conception and design: NM, SVK

Acquisition of experimental data: NM

Modeling: NM, SS, PWW, SVK

Analysis and interpretation of data: NM, SVK

Drafting of Manuscript: NM, SVK

All authors contributed to the critical revision and approval of the final manuscript.

## Acknowledgements

This work was supported by Natural Sciences and Engineering Research Council (NSERC, RGPIN-288253 and RGPIN-2017-05005), Canadian Institutes for Health Research (CIHR MOP-77643). NM was supported by the Faculty of Dentistry McGill University and le Réseau de Recherche en Santé Buccodentaire et Osseuse (RSBO). Special thanks to Dr. M. Murshed (McGill, Montreal) for C2-OB cells and Dr. P. Grutter and his graduate students M. Anthonisen and M. Rigby (McGill, Montreal) for glass capillary puller.

